# Leaf shape modulates climate–trait relationships in the wild species *Chenopodium hircinum* (Amaranthaceae)

**DOI:** 10.1101/2025.04.21.649804

**Authors:** Jonatan Rodríguez, Vilma Quipildor, Eugenia Giamminola, Sergio J. Bramardi, David Jarvis, Jeff Maughan, Jiemeng Xu, Hafiz U. Farooq, Pablo Ortega-Baes, Eric Jellen, Mark Tester, Daniel Bertero, Ramiro N. Curti

**Affiliations:** Laboratorio de Investigaciones Botánicas (LABIBO), Facultad de Ciencias Naturales, Universidad Nacional de Salta, Av. Bolivia 5150, Salta, Argentina; Consejo Nacional de Investigaciones Científicas y Técnicas (CONICET) CCT Salta-Jujuy, Argentina; Centro de Investigaciones en Toxicología Ambiental y Agrobiotecnología del Comahue (CITAAC-CONICET), Universidad Nacional del Comahue, Neuquén, Argentina; Department of Plant and Wildlife Sciences, Brigham Young University (BYU), Provo, UT, USA; Biological and Environmental Sciences and Engineering Division, King Abdullah University of Science and Technology (KAUST), Thuwal, Saudi Arabia; Facultad de Agronomía, Universidad de Buenos Aires and Instituto de Investigaciones Fisiológicas y Ecológicas Vinculadas a la Agricultura (IFEVA-CONICET), Buenos Aires, Argentina

## Abstract

Understanding how leaf morphology mediates plant responses to environmental variation is essential for predicting species adaptability under climate change. In this study, we investigated natural variation in leaf shape and associated functional– physiological traits (FPTs) across populations of *Chenopodium hircinum* grown in a common garden. We found that leaf shape strongly correlates with the climatic conditions of population provenance, while functional and physiological traits are independently associated with morphology rather than directly with climate. Landmark-based morphometric analysis revealed that the second principal component, distinguishing deeply lobed from rounded leaves, is significantly linked to key traits such as leaf mass per unit area (LMA) and stomatal conductance. These relationships suggest that morphology mediates a functional continuum between resource-use strategies. Notably, within-population phenotypic variation accounted for much of the observed trait diversity, underscoring the role of individual-level phenotypic plasticity in shaping ecological function. Our results challenge traditional views on the thermoregulatory role of leaf lobation and highlight morphology as a central axis of adaptation. This work advances understanding of trait integration in response to environmental heterogeneity and suggests that plasticity in leaf shape and function may enhance resilience of *C*. *hircinum* across its native range.

## Introduction

The functional significance of leaf shape remains a central topic in plant evolutionary ecology and ecophysiology (Niklas et al. 2023). The extraordinary diversity of leaf morphology among terrestrial plants—both across species and within individuals— represents a key evolutionary innovation (Ferris 2019; Nakayama 2024). As primary organs of carbon assimilation, leaves are subject to strong selective pressures (Niklas 1999), shaping ecological strategies and evolutionary pathways (Niinemets 2020). This morphological variation facilitates plant adaptation to diverse habitats, particularly given the sessile nature of plants and the leaf’s critical role as the interface with the surrounding environment (Vogel 2009).

Recent studies of herbaceous species demonstrate that environmental variables—especially temperature—exert a strong influence on leaf shape at both interspecific and intraspecific levels (Mason and Donovan 2015; Giupponi 2020; Hightower et al. 2024; Singhal et al. 2025; Curti et al. 2025), mirroring patterns observed in other plant groups (Peppe et al. 2011; Royer et al. 2012; McKee et al. 2019). Several of these investigations link morphological variation to climatic gradients and adaptive functional traits. For instance, in *Helianthus*, leaf shape was associated with traits supporting climatic adaptation at both inter- and intraspecific scales (Mason and Donovan 2015). Likewise, Giupponi (2020) found that intraspecific leaf shape variation in *Campanula elatinoides* aligned with traits indicative of climatic adaptation. Most recently, Singhal et al. (2025) reported a latitudinal gradient in leaf shape linked to thermoregulatory capacity in *Ipomoea hederacea*.

Prior studies exploring the relationship between leaf morphology and environment have typically contrasted broad categories of leaf types—e.g., entire versus dentate or lobed versus unlobed—and related these to physiological functions such as thermoregulation and water balance (Royer and Wilf 2006; Nicotra et al. 2011). Entire, large-surfaced leaves have often been linked with efficient transpirational cooling in warm climates (Nicotra et al. 2011; Singhal et al. 2025). Nonetheless, physiological trade-offs—such as differences in chlorophyll content—remain largely species-specific (Mason and Donovan 2015; Everingham et al. 2024), and temperature appears to exert limited effects on leaf mass per area (Niklas et al. 2023). Importantly, many herbaceous taxa exhibit substantial intraspecific variation in leaf shape that goes beyond binary classifications. In particular, lobed-leaf species may show variation in the number, depth, and position of lobes, as well as in their distribution along the lamina (Gupta et al. 2020; Balant et al. 2024; Hightower et al. 2024). This raises the question: can such subtle within-species variation in marginal complexity contribute to functional traits involved in thermal adaptation? Focused intraspecific analyses are needed to clarify the extent to which fine-scale morphological differences mediate physiological responses across environmental gradients.

A recent study assessed the utility of various leaf shape descriptors—including geometric morphometric approaches such as landmark- and outline-based analyses—in capturing morphological variation in *Chenopodium hircinum* (Curti et al. 2025), the wild progenitor of quinoa (*C*. *quinoa*) (Jarvis et al. 2017; Rey et al. 2023; Maughan et al. 2024). Native to South America, *C*. *hircinum* is primarily distributed in Argentina (Wilson 1990; Curti et al. 2023), where it is increasingly studied for traits that may confer climate resilience to quinoa (Curti et al. 2022; Agüero-Martínez et al. 2024; Curti et al. 2025; Xu et al. 2025). The species occurs across a broad elevational and thermal gradient in Argentina (Curti et al. 2023), and recent findings suggest that morpho-phenological and leaf-shape variation are shaped more by individual and within-population differences than by broad ecoregional patterns (Curti et al. 2022; Curti et al. 2025). Moreover, trait variation correlates with the east–west temperature gradient, suggesting a high degree of phenotypic integration (Curti et al. 2022; Curti et al. 2025).

In Argentinean populations of *C*. *hircinum*, geometric morphometric approaches outperformed traditional shape metrics (e.g., aspect ratio, circularity, solidity) in capturing leaf shape variation. Landmark and elliptical Fourier descriptors revealed clear population-level differentiation in lobulation patterns along the blade: basal lobulation predominates in high-elevation, colder sites, whereas mid-blade lobulation is more common in warmer, low-elevation populations (Curti et al. 2025). If such thermal gradients influence leaf morphology, populations across these gradients should also differ in physiological and functional traits. In addition, given the high morphological and physiological plasticity of ruderal species in general (Everingham et al. 2024; Hočevar et al. 2025), and of *C*. *hircinum* in particular (Curti et al. 2022; Curti et al. 2025), we hypothesize that leaf shape will more accurately predict physiological than functional trait variation. In this study, we address the following questions: Does variation in leaf lobulation influence physiological and functional traits independently? If so, is this influence primarily driven by species-level or individual-level variation?

## Materials and Methods

### Collection sites, common garden experiment and leaf sampling

This study analyzed seeds from 11 populations of *Chenopodium hircinum* collected within the Salta and Jujuy provinces of Northwest Argentina (Table S1). The majority of seeds originated from a recent collection undertaken during January and February of 2023, while seeds from two populations (LSK and SCA) were sourced from a collection in January 2022. The ecoregions represented by these seed origins are High Monte, Dry Chaco, and Central Andean Puna, in descending order of sample representation (Table S1). Mean summer temperature (MST, °C) and accumulated summer precipitation (ASP, mm) data for the collection sites during the austral summer months (January to March) were acquired from WorldClim (https://www.worldclim.org/) at a spatial resolution of 30 arc-seconds (approximately 1 km² at the equator).

A common garden experiment was conducted ex situ between January and March of 2024 at the Experimental Field of the National University of Salta (24.72° S, 65.41° W, 1228 m a.s.l.) under outdoor conditions. Seeds underwent a dormancy-breaking protocol as described by Curti et al. (2022). Following germination in Petri dishes (indicated by radicle emergence), seedlings were transplanted into 7-liter pots arranged in a completely randomized design, with ten replicate pots per population. Pots were filled with a 3:1 (v/v) mixture of sand and peat substrate, initially sown with five plants per pot and subsequently thinned to a single plant. To mitigate potential edge effects, pots were randomly repositioned every two days.

A total of 53 fully expanded leaves were sampled at the anthesis stage from three to eight individual plants per population (Table S1). Specifically, leaves were harvested from the middle third of the main stem of each plant, coinciding with the location where physiological and functional traits were also assessed (see below). After petiole excision, leaves were positioned adaxially on graph paper and photographed using a digital camera (Nikon D7200, Nikon Corporation Hong Kong) at a consistent distance. Images were saved as standard JPEG files for subsequent image analysis.

### Evaluation of leaf shape, physiological and functional traits

Shape descriptors, including circularity (CIRC), aspect ratio (AR), and solidity, were computed using ImageJ (https://imagej.net/ij/) following the conversion of leaf images to binary silhouettes (Curti et al. 2025). Eight previously established landmarks on the leaf outline of *C*. *hircinum* were digitized using the ImageJ point tool (Curti et al. 2025). Additionally, Elliptical Fourier Descriptors (EFDs) were calculated according to the methodology outlined in Curti et al. (2025).

Three physiological traits were measured on clear sunny days between 11:00 am and 16:00 pm on previously irrigated pots at dawn. Measurements included stomatal conductance (*g_s_*, umol H_2_O m^-2^ s^-1^) using a leaf porometer (SC-1 Meter Group USA 2021), chlorophyll concentration (CHL, umol m^-2^) using a MC-100 (Apogee Instruments INC, USA), and leaf temperature (°C, LTP) with an infrared thermometer (GM320, China). Three functional traits were evaluated in the same leaves that were used for physiological measurements. Leaf dry mass per unit area (mg mm^-2^, LMA) was determined by weighting leaves (discarding the petiole, LDW mg) and computing leaf area (mm^-2^, LA) from digitalized leaf image.

### Data analysis

All statistical analyses were performed within the R environment (R Core Team, 2024). Initial analyses involved a general description of leaf shape based on landmarks and leaf outlines, following a previously published protocol (Curti et al. 2025). Briefly, landmark coordinates were processed using the ‘shapes’ package (Dryden and Mardia 2016) after undergoing Generalized Procrustes Analysis (GPA). Subsequently, Principal Component Analysis (PCA) was applied to the Procrustes-aligned coordinates, and Principal Component (PC) scores for all retained components (hereafter, Ldk PCs) were determined using the broken-stick model implemented in the ‘PCDimension’ package (Wang et al. 2017). The analysis of leaf outlines followed a similar procedure: coordinates of leaf silhouettes were acquired using the ‘Momocs’ package (Bonhomme et al. 2014), and outlines were normalized via a full GPA adjustment based on the eight previously defined landmarks. Following alignment, Elliptical Fourier Transforms were fitted separately for the *x* and *y* coordinates. The number of harmonics to be retained for subsequent analysis was estimated based on accumulated harmonic power (Bonhomme et al. 2014), and PC scores for the retained components (hereafter, EFD PCs) were extracted using the ‘PCDimension’ package.

The relationships among Functional and Physiological Traits (FPTs: LDW, LMA, LA *g_s_*, LTP, and CHL), shape descriptors (CIRC, AR, solidity, retained Ldk PCs, and EFD PCs), and climatic conditions (Elevation, MST, and ASP) were assessed using Escoufier’s RV coefficients (‘FactoMineR’ package, (Josse et al. 2008; Lê et al. 2008)) with standardized variables. A Multivariate Multiple Regression (MMR) analysis (‘car’ package, (Fox et al. 2001)) was employed to test whether shape and climatic variables significantly predicted FPTs, utilizing a MANOVA (Pillai’s trace statistic). Subsequently, multiple linear regression (MLR) analyses were fitted to identify which predictor variables accounted for variation in each FPT individually. A second MMR analysis was conducted using significant predictor variables identified in the initial MLR analyses, again employing MANOVA with the Pillai statistic. Correspondingly, MLR analyses were performed to further examine the influence of these significant predictor variables on individual FPTs. A comparison between the full model (first MMR analysis) and the simplified model (second MMR analysis) was performed using an ANOVA. Finally, Redundancy Analysis (RDA) was performed using the ‘vegan’ package (Oksanen et al. 2001) to visualize the results of the best-fitting MMR model. The significance of the model, predictors (shape and climatic variables), and the number of RDA axes required to explain variation in FPTs were evaluated using a permutation test. A triplot displaying the relationships among the matrices of FPTs, samples, and predictors on the retained RDA axes was constructed for visual interpretation.

## Results

### General patterns of leaf shape variation

From the 16 Ldk PCs obtained, only the first two (Ldk PC1 and Ldk PC2) were important to capture significant variation in leaf shape among samples (Fig. 1a). The Ldk PC1 was associated with variations in the relative distances between landmarks 3 and 4, and 6 and 7, distinguishing leaves with pronounced lobulation in the middle section from those with less lobulation in this region (Fig. 1a). In turn, the Ldk PC2 differentiated samples with deeply lobed leaves (trilobed) from those with rounded leaves mainly associated with variation in landmarks 1, 2, 3, 4, 6, and 7 (all of them at the central position along the leaf blade) (Fig. 1a).

**Fig. 1.**
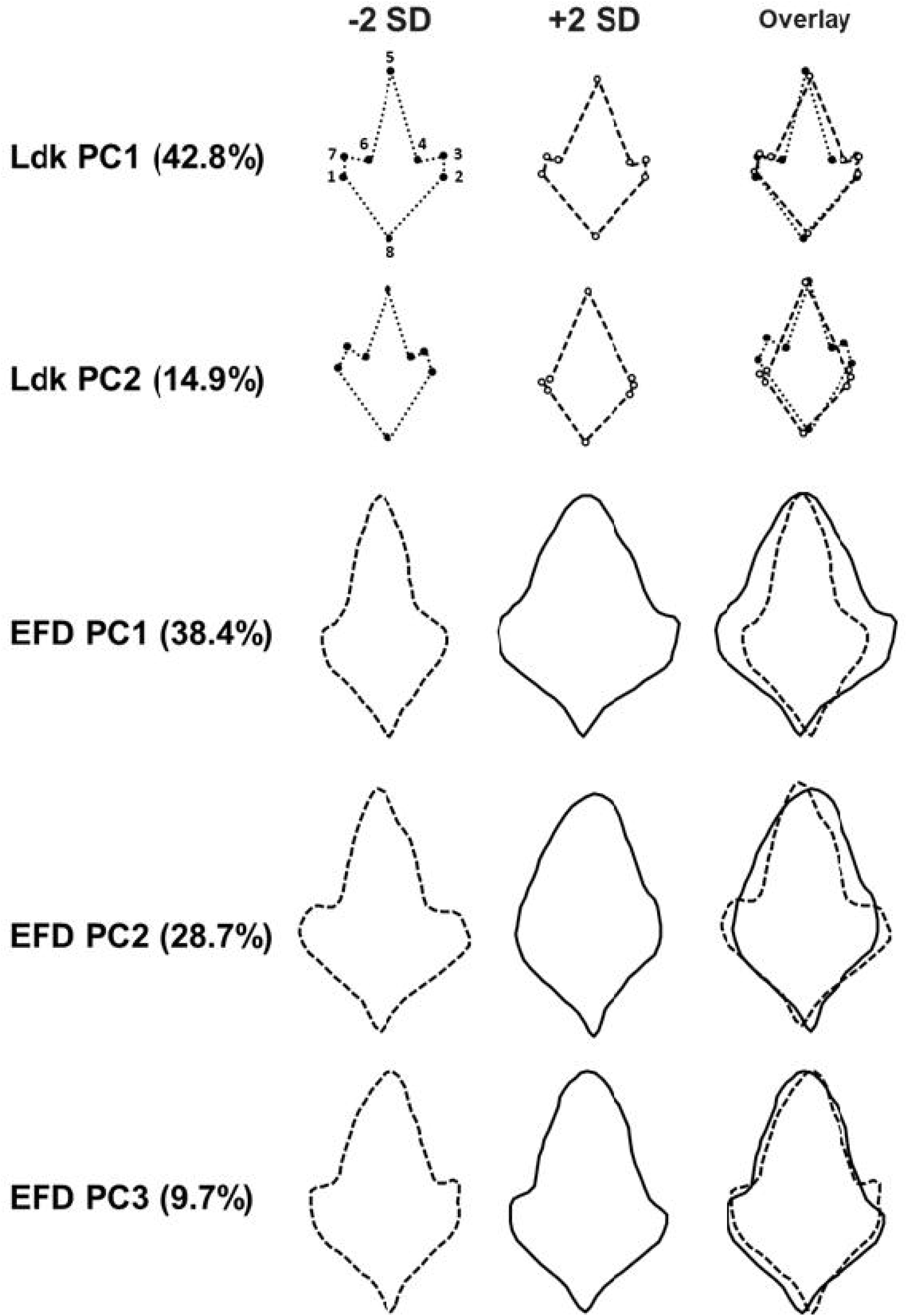
Principal components analyses (PCA) were performed using both landmark coordinates and Elliptical Fourier Descriptors (EFDs) to generate representative “eigenleaves” summarizing shape variation. For each principal component (PC) that significantly contributed to morphological variation, the proportion of explained variance is reported. Shape changes along each PC axis are visualized using reconstructed leaf outlines at ±2 standard deviations (SD) from the mean—solid lines for −2 SD and dashed lines for +2 SD—demonstrating the extent of variation captured. Key shape differences are further emphasized by overlaying these opposing outlines. The positions of eight key morphological landmarks are also indicated.

Fourteen harmonics were necessary to describe leaf outlines (results not shown), whereas only the first three Elliptical Fourier Descriptors PCs captured significant leaf shape variation among samples (Fig. 1b). The EFD PC1 reflects overall leaf outline variations, distinguishing samples with smoother and uniform contours (left) from samples with irregular and wider shapes with an expanded base (Fig. 1b). The EFD PC2 captures changes in leaf area within the middle third, associated with an enlargement of mild-lobing development, differentiating samples with deeply lobulation (trilobed) from samples with rounded leaves (Fig. 1b). On the other hand, EFD PC3 was associated with the development of lobes at the right side in middle-third along leaf blade (Fig. 1b).

### Interrelationships between shape, climate and FPTs

A significant association was found between functional–physiological traits (FPTs) and shape descriptors, as well as between shape and climate matrices; however, FPTs showed no direct relationship with climate variables (Table 1a). A follow-up analysis, separating functional and physiological traits and examining each category of shape descriptors, yielded consistent results. Functional traits were significantly associated with all shape descriptors, whereas physiological traits correlated only with landmark- and Fourier-based measures (Table 1b). Notably, functional and physiological traits were not interrelated. In contrast, all shape descriptors showed significant associations with climate factors and with each other (Table 1b).

**Table 1.**
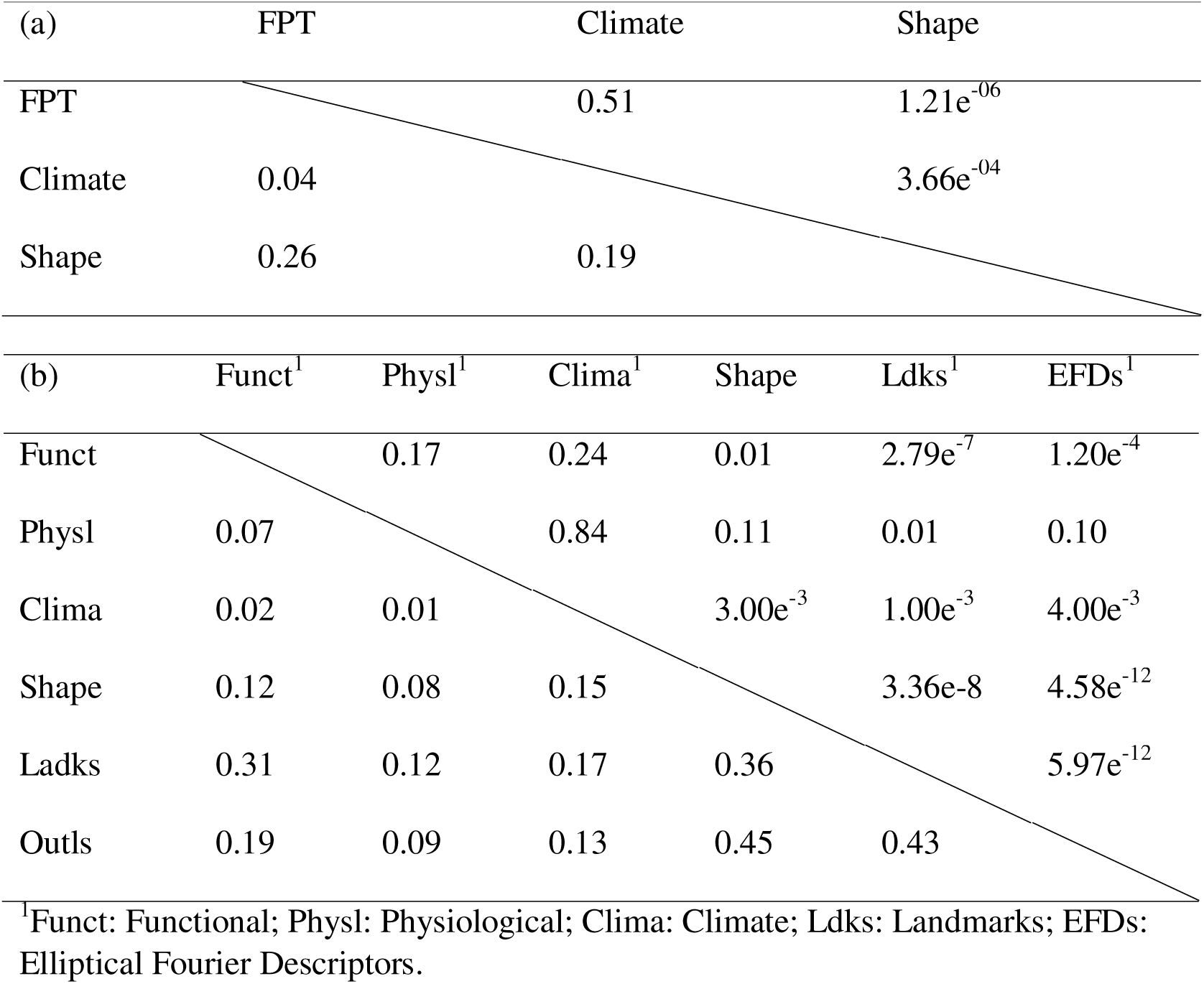
RV coefficients (below diagonal) and their significance (*p*-values, upper diagonal) for the associations between Functional and Physiological Traits (FPTs), Climate and Shape descriptors matrices on a general basis (a) and an individual basis (b).

The full model of MMR analysis revealed that shape descriptors and climate factors were significant to predict leaf FPTs (Pillai test statistics = 1.987, approx. *F* _(66, 246)_ = 1.846, *p* = 0.00042). The significant predictors variables were mainly Ldk PC2 for most traits, and MST and aspect ratio plus Ldk PC2 for LTP (Fig. 2b, c, d, e). None predictor variables explained variation in both traits LDW and CHL (Fig. 2a, b). The second MMR analysis considering only MST, aspect ratio and Ldk PC2 predictors variables was also significant (Pillai test statistics = 0.764, approx. *F* _(18,138)_ = 2.622, *p* = 0.00083). The significant predictors variables were mainly aspect ratio and Ldk PC2 for most traits (Fig. 2a, b, c, d, e, f). The comparison between full and reduced models was significant (Pillai test statistics = 1.41, approx. *F* _(48, 246)_ = 1.574, *p* = 0.014), implying that the reduced model considering only significant predictors was better to describe variation in FPT.

**Fig. 2.**
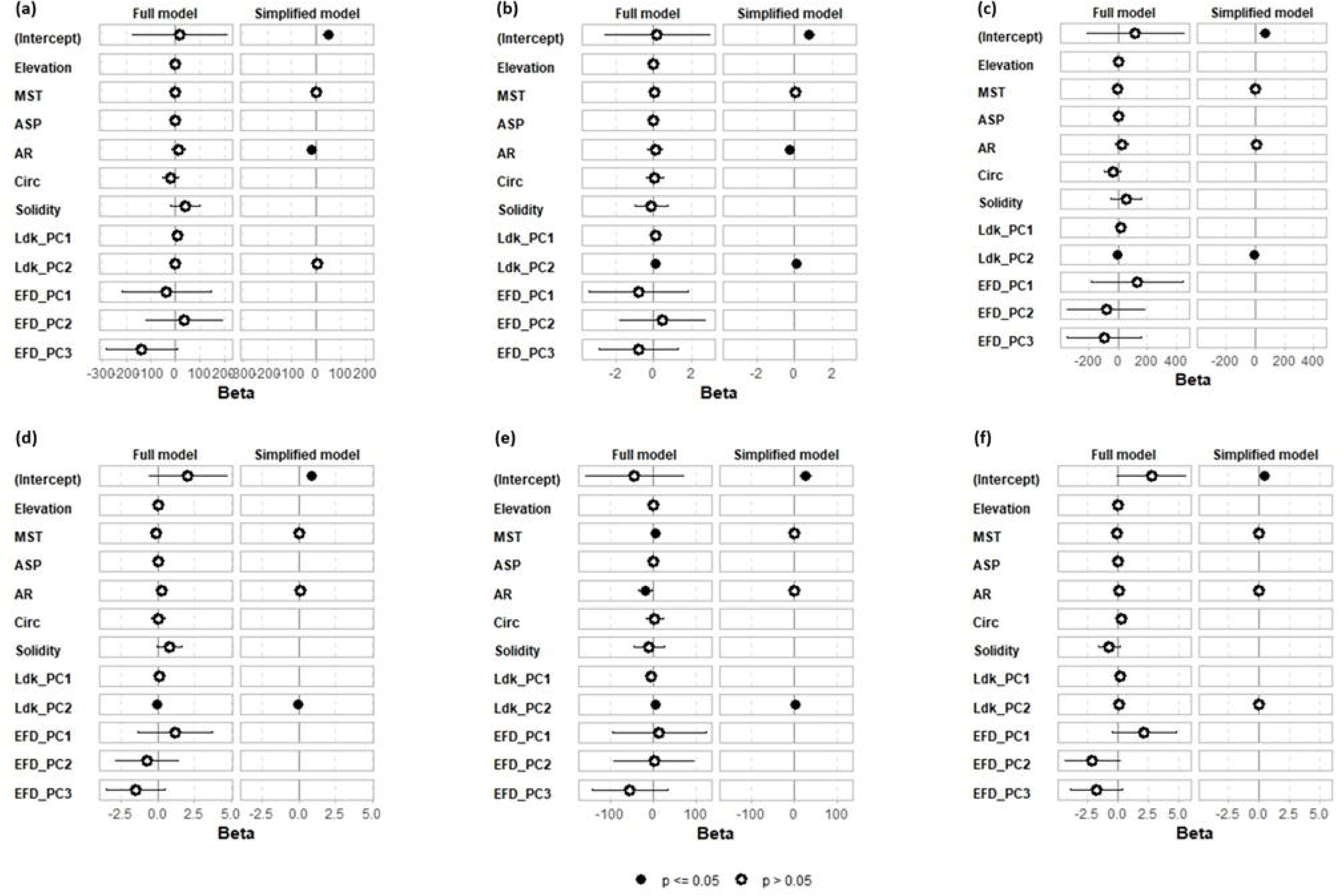
Plots illustrating the coefficients of both the full and simplified models for (a) LDW, (b) LA, (c) LMA, (d) *g_s_*, (e) LTP, and (f) CHL.

The Redundancy Analysis (RDA) showed that shape and climatic variables explain 21% of the variation in FPTs responses. The permutation test confirmed the significance of the MMR reduced model, with only shape descriptors (aspect ratio and Ldk PC2) contributing to FPTs variation (Table S2). Only the first RDA component was significant (Table S2). In the triplot, LMA and *g_s_* were positively associated with aspect ratio and positioned above the origin, while LA and LTP were linked to Ldk PC2 and positioned below (Fig. 3). Samples distributed along the first RDA axis showed strong admixture among populations (Fig. 3). CHL and MST were near the origin, aligning with their low significance and poor predictive capacity for FPTs responses.

**Fig. 3.**
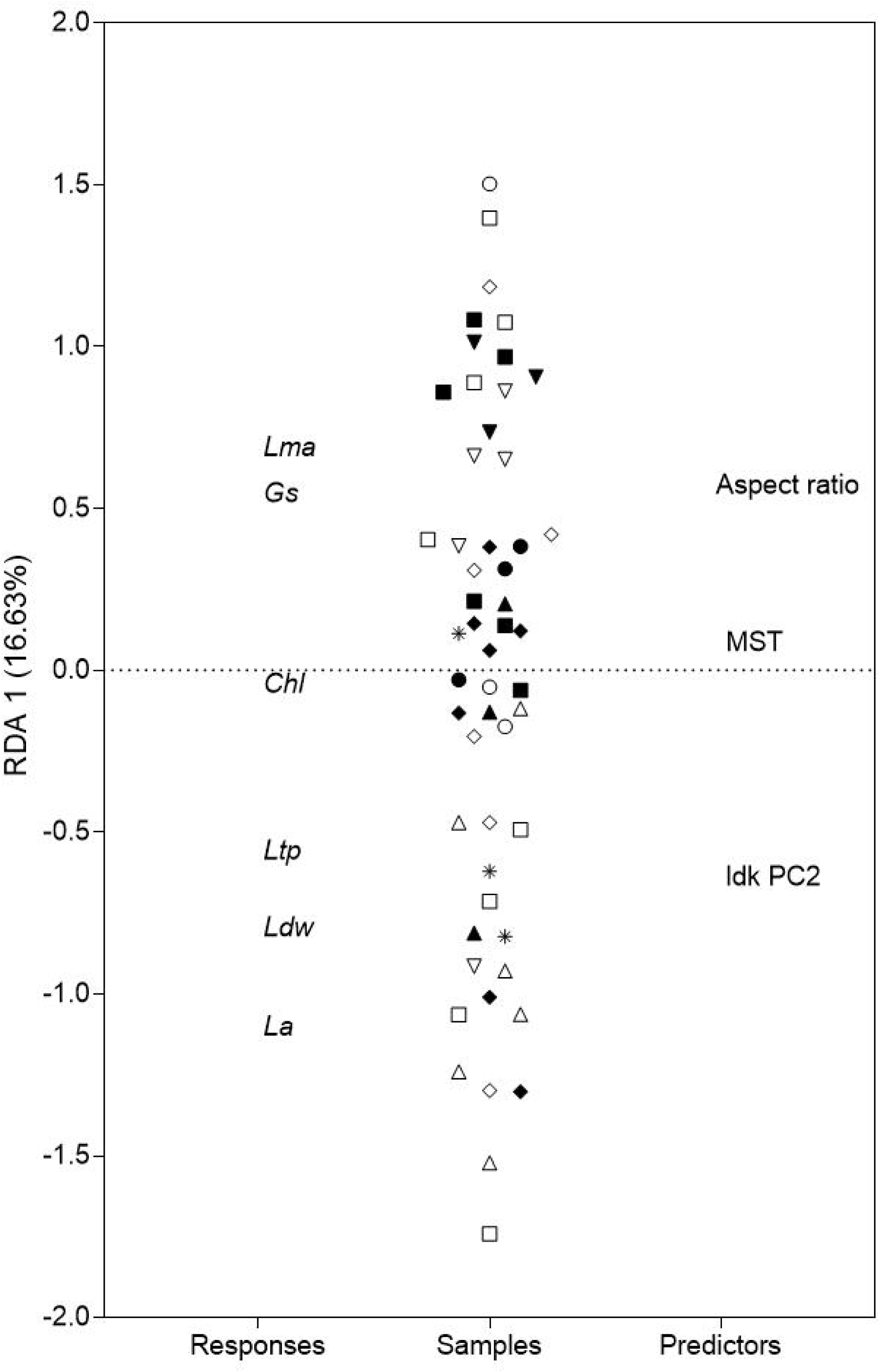
Triplot of the 1st component of RDA of *C*. *hircinum* samples, functional-physiological traits (responses) and attributes (predictors). Populations are displayed by symbols.

## Discussion

The principal findings of this study are threefold: (1) leaf shape variation among *C*. *hircinum* populations correlates strongly with the climatic conditions of their provenance sites; (2) physiological and functional traits are independently associated with leaf morphology; and (3) a substantial proportion of field-observed variation arises from within-population phenotypic plasticity. These results underscore a robust association between leaf form, physiological–functional attributes, and local climate, suggesting that environmental heterogeneity is a key driver of the ecological relevance of leaf morphology in *Chenopodium hircinum*. Furthermore, the marked contribution of within-population phenotypic plasticity highlights a critical mechanism through which *C*. *hircinum* adapts to environmental variability.

The relationships among leaf shape, functional–physiological traits (FPTs), and climate reveal a morphology-mediated pathway linking climate to plant function. While significant correlations emerged between shape descriptors and both FPTs and climatic variables (Table 1a), direct associations between climate and either functional or physiological traits were absent (Table 1b). Additionally, although functional and physiological traits were each strongly associated with shape, they were not significantly correlated with one another (Table 1b). This pattern indicates that leaf morphology acts as an independent mediator between environmental factors and physiological function. Landmark-based analyses identified the second principal component (PC2) as the only axis significantly distinguishing deeply trilobed from rounded leaves (Fig. 1), with PC2 showing strong correlations with multiple FPTs (Table 1b), as evidenced by multivariate multiple regression (MMR) and multiple linear regression (MLR) analyses (Fig. 2).

Redundancy Analysis (RDA) of the best-fit MMR model revealed a significant co-localization of leaf mass per unit area (LMA) and stomatal conductance (gs) (Fig. 3). Elevated *g_s_* was positively correlated with aspect ratio and negatively correlated with both leaf temperature and lobation (i.e., lower PC2 scores), consistent with theoretical expectations that lanceolate, entire-margined leaves operate at sub-ambient temperatures via high transpiration (Nicotra et al. 2011; Singhal et al. 2025). However, these results also challenge the hypothesized thermoregulatory role of lobation, as lobed leaves exhibited higher temperatures despite comparable area (Fig. 3).

The spatial spread of samples along PC2 further showed that lobed leaves extended beyond the morphological range of rounded ones (Fig. 1), complicating efforts to test whether lobation reduces internal heat transfer distances (Nicotra et al. 2011). Moreover, the observed positive correlation between gs and LMA, coupled with a negative correlation between leaf dry weight (LDW) and LMA (Fig. 3), suggests a resource-acquisitive strategy in *C*. *hircinum*. This pattern aligns with broad global trends (Wright et al. 2004) and mirrors findings in Helianthus species, where enhanced gas exchange correlates with leaf shape at both intra- and interspecific levels (Mason and Donovan 2015). Notably, our findings highlight a clear link between this resource-use strategy and contrasting leaf shape configurations—an association that remains underreported in prior studies of leaf form–function relationships.

Deeply lobed leaves (low PC2 scores) were also larger in area (LA) and had higher dry mass (Fig. 3). An allometric analysis revealed a slope < 1 between these traits (Fig. 4), consistent with the concept of diminishing returns in leaf mass investment (Niklas et al. 2007). Yet, tissue bulk density (LMA)—a proxy for lamina stiffness—was inversely related to LDW, suggesting it does not fully explain the increase in leaf mass. This points to additional structural factors such as leaf thickness as potential contributors to elevated LMA, a hypothesis meriting further investigation in *C*. *hircinum* (Shi et al. 2024). This observation supports the idea that individual components of complex traits can vary independently and respond at different rates to environmental change (Niinemets 1999; Poorter et al. 2009).

**Fig. 4.**
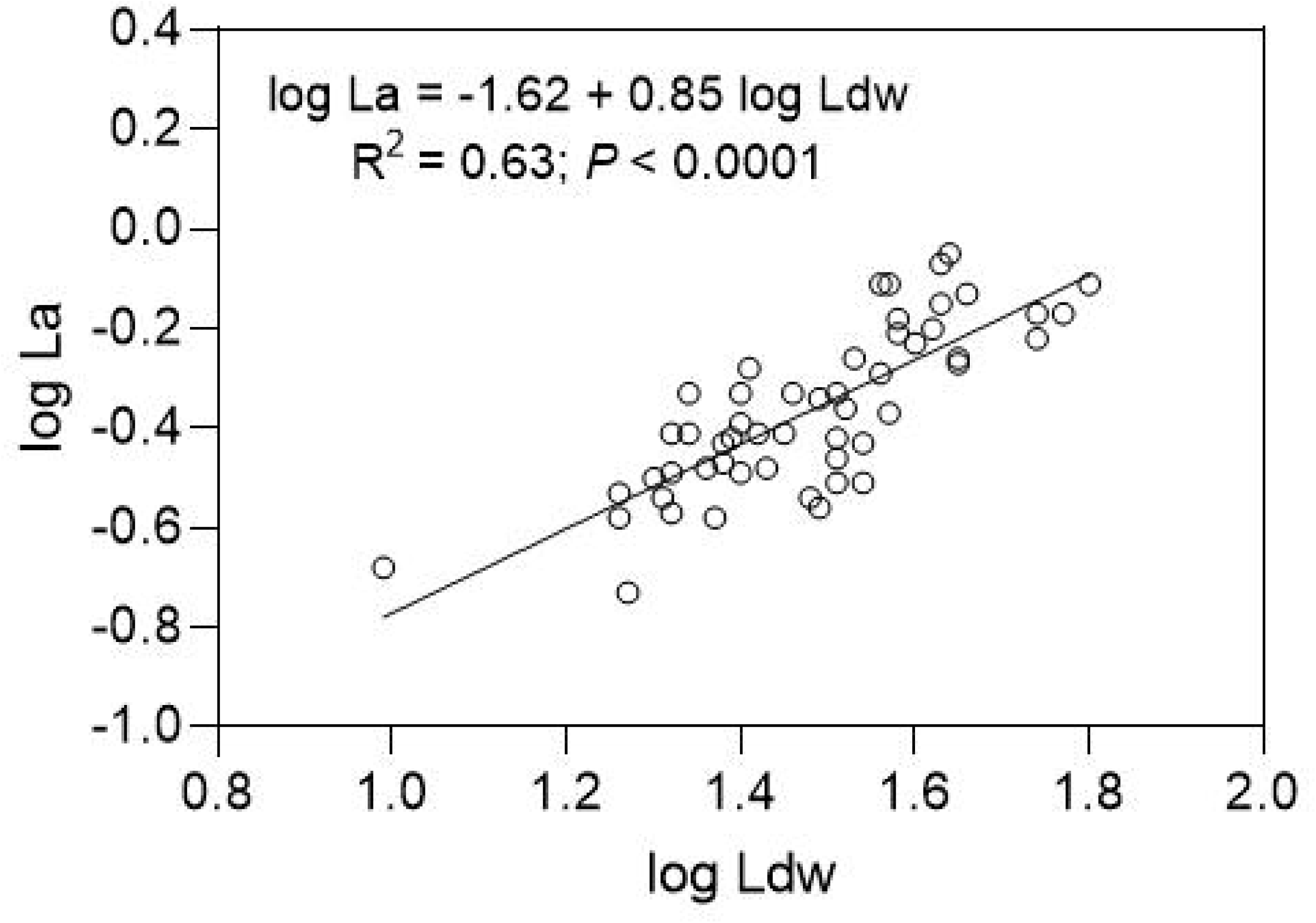
Log–log bivariate relationship for LA vs. LDW. Information about the regression model, coefficient of determination (*R*^2^) and significance (p-value) are shown.

A key insight from this study is the broad range of phenotypic variation in FPTs observed within *C*. *hircinum* populations grown under common garden conditions. As phenotypic variation under controlled environments does not inherently reflect environmental influences (Niinemets 2020), future experiments should explore trait expression across contrasting environments—e.g., across growing seasons or via reciprocal transplants. Such approaches could help disentangle the roles of genetic variation and plasticity in driving the observed trait diversity.

Our results indicate that functional and physiological traits varied independently, yet exhibited complex interrelationships (Fig. 3). This is particularly relevant given the broad environmental heterogeneity characterizing the species’ native range (Curti et al. 2022, 2023, 2025). Importantly, our findings emphasize the individual-level nature of trait variation, rather than strict ecotypic differentiation, paralleling results in other wild species (Westerband et al. 2021; Chandler and Travers 2024). In light of climate change, intra-population phenotypic plasticity emerges as a crucial mechanism by which *C*. *hircinum* may track rapid environmental shifts. Our study identifies potential pathways—particularly those mediated by leaf morphology—through which this species may respond to changing climatic conditions across its geographic range.

## Conclusions

This study reveals a robust, morphology-mediated linkage between climate, functional– physiological traits (FPTs), and leaf shape in *Chenopodium hircinum*. Leaf morphology emerged as a central integrator, independently connecting climatic variables with physiological and functional trait expression. Notably, deeply lobed and rounded leaf forms corresponded to distinct trait syndromes, suggesting divergent ecological strategies within populations. Furthermore, the pronounced within-population variation observed under common garden conditions points to a substantial role for phenotypic plasticity in shaping trait expression. This plasticity, particularly at the individual level, may serve as a key mechanism for adaptive responsiveness across heterogeneous environments. In the context of rapid climate change, such intra-population plasticity could facilitate persistence and functional versatility across *C*. *hircinum*’s geographic range. Collectively, these findings underscore the ecological significance of leaf shape beyond its taxonomic utility and emphasize the importance of integrating morphological analyses into studies of trait–environment interactions. Future research incorporating experimental manipulations and multi-environment trials will be essential for disentangling the relative contributions of plasticity and genetic differentiation to trait variation in this widespread species.

## Supplementary data

Table S1. Passport data of *Chenopodium hircinum* populations evaluated in the present study

Table S2. Redundancy analysis (RDA) results showing the significance of the simplified MMR model, associated climate factors and shape descriptors, and retained axes

## Data availability

The data that support this study will be shared upon reasonable request to the corresponding author.

