## Supplementary material for "Leaf shape modulates climate–trait relationships in the wild species *Chenopodium hircinum* (Amaranthaceae)": Table S1 and Table S2

Table S1. Passport data of *Chenopodium hircinum* populations evaluated in the present study

| Pop. | Ecoregion | Province | Department | Site | Lat. (S) | Lon. (W) | Elevation (masl) | Samples (*n*) | MST (°C) | ASP (mm) |
| --- | --- | --- | --- | --- | --- | --- | --- | --- | --- | --- |
| H17 | Dry Chaco | Salta | Rosario de la Frontera | Rosario de la Frontera | 25.81 | 64.98 | 792 | 3 | 22.87 | 416 |
| H36 | Dry Chaco | Jujuy | El Carmen | Los Lapachos | 24.48 | 65.07 | 806 | 5 | 23.37 | 345 |
| H05 | Dry Chaco | Salta | La Viña | Ampascachi | 25.36 | 65.53 | 1158 | 3 | 21.70 | 271 |
| SCA | High Monte | Salta | San Carlos | San Carlos | 25.88 | 65.93 | 1633 | 3 | 21.13 | 115 |
| H30 | High Monte | Salta | San Carlos | Las Viñas | 25.80 | 65.96 | 1688 | 3 | 20.97 | 108 |
| LSK | High Monte | Salta | San Carlos | Las Viñas | 25.79 | 65.96 | 1738 | 6 | 20.73 | 102 |
| H28 | High Monte | Salta | San Carlos | Payogastilla | 25.70 | 66.01 | 1766 | 6 | 20.37 | 105 |
| H25 | High Monte | Salta | San Carlos | Angostura | 25.51 | 66.23 | 1977 | 6 | 19.50 | 96 |
| H22 | High Monte | Salta | Cachi | Cachi | 25.10 | 66.19 | 2477 | 8 | 17.17 | 112 |
| RI5 | High Monte | Jujuy | Humahuaca | San Roque | 23.26 | 65.36 | 2898 | 7 | 16.00 | 121 |
| H01 | Central Andean Puna | Salta | Rosario de Lerma | Santa Rosa de Tastil | 24.45 | 65.95 | 3119 | 3 | 14.43 | 113 |

MST: mean summer temperature (January to March, austral summer); ASP: accumulated summer precipitation.

Table S2. Redundancy analysis (RDA) results showing the significance of the reduced MMR model, associated climate factors and shape descriptors, and retained axes.

|  | MS | *F-value* | *P-value* |
| --- | --- | --- | --- |
| Model | 1.27 | 4.37 | 0.001 |
| MST | 0.06 | 0.58 | 0.71 |
| Aspect ratio | 0.57 | 4.94 | 0.001 |
| ldk_PC2 | 0.73 | 7.59 | 0.001 |
| RDA1 | 0.99 | 10.32 | 0.001 |
| RDA2 | 0.25 | 2.58 | 0.08 |
| RDA3 | 0.02 | 0.19 | 0.96 |
| Residual | 4.73 |  |  |
